# Peers over the past: Prior predation-risk experience does not dictate antipredator responses of individuals in groups

**DOI:** 10.1101/2024.12.31.630853

**Authors:** Kanika Rawat, Kavita Isvaran

**Author notes:** Corresponding Author Kanika Rawat, TB-02, Centre for Ecological Sciences Indian Institute of Science, Bengaluru, Karnataka India-560012.

## Abstract

Animals with predation-risk experiences can use the information to modulate their antipredator responses and survive future risk conditions (behavioural carryover). However, different contexts, such as sociality, may alter the risks and associated responses for an individual. A group can provide benefits to an individual, such as dilution of risk, confusion effect, and improved vigilance, potentially modifying carryover payoffs. Therefore, we tested the impact of sociality on the behavioural carryover of predation-risk experience. We used *Aedes aegypti* as a model system, where we had already established carryover, showing that predation-risk experience dictates future behaviour in a solitary setting. In contrast to solitary settings, we predicted that the behavioural carryover would not manifest in a group setting due to differences in trade-offs. We compared the behaviour of experienced and naive individuals in group settings in the presence and absence of an immediate threat. Our findings revealed that the past predation-risk experience did not influence the behaviour of individuals in a group setting. Instead, the immediate threat environment affected experienced and naive individuals similarly. These results suggest that the experience from predation past is more relevant when prey are solitary than when they are in a group due to the additional group advantages.

## Introduction

Behavioural carryover of risk experiences is a valuable antipredator tactic. Prey use their prior experiences to improve their response against a potential threat, and such carryovers can occur across an organism’s life cycle or generations (Brass et al., 2020; Garcia et al., 2019; West et al., 2018). Similar to any costly antipredator behaviour, behavioural carryovers of predation-risk experience may manifest based on the needs and payoffs across different contexts that can vary the vulnerability of prey and influence the trade-offs between different fitness components (Brown & Godin, 2023; Wirsing et al., 2021). However, the altered manifestations of behavioural carryovers under ecologically relevant contexts remain understudied.

Sociality or group living can heavily influence the vulnerability of prey. While group living has its costs (Ezenwa et al., 2016; Landry & Li, 2022), the benefits of being in a group outweigh them in the face of predation (Rubenstein, 1978). A group can benefit an individual through shared vigilance, social information transfer, confusion and dilution effects (Lehtonen & Jaatinen, 2016), decreasing the predation risk for an individual in a group. Therefore, the expression of antipredator behaviour varies for an individual from solitary to group settings. For example, individual sun skinks decrease the time allotted to antipredator behaviour as the group size increases in high-risk conditions (Downes & Hoefer, 2004). Similarly, female Seychelles warblers make alarm calls more frequently when they encounter a nest predator model in the absence of their partners but will actively attack the model when with their partners (Groenewoud et al., 2019). Sociality not only alters individual antipredator behaviour but also facilitates the emergence of collective antipredator strategies. These collective behaviours, such as mobbing predators (Cunha et al., 2017) or using confusing collective movements to escape attacks (Couzin & Krause, 2003), arise from the coordinated actions of group members.

With predation being a strong selective force, animals exhibit multiple traits to avoid the risk of predation. However, the interplay and emergent outcomes of these various traits remain poorly understood. We know very little about the influence of prior predation-risk experience on an individual’s behaviour when it is in a group setting. Since being in a group is an effective antipredator strategy, using prior predation-risk experience to improve an individual’s response may be redundant in a group setting, or it may further add to the advantages by enhancing the group response. A recent study on common cyprinid fish shows a starker difference in the activity of predator-naive and experienced fish when solitary than in a group setting. The authors suggested that this trend could be due to decreased predation risk, increased competition for food, or other factors, all of which warrant further investigation (Wang et al., 2019).

To investigate the behavioural consequences of predation-risk experiences in a group, we use the pupal stage of *Aedes aegypti* as the model system. The pupal stage is immature and does not feed; the absence of competition for food strikingly lowers the cost of being in a group and reduces the confounding factors potentially affecting behaviours of interest, such as activity. Consequently, we can gain a deep insight into the functional roles of behaviours of interest against a background of fewer interacting selection pressures. Therefore, the mosquito pupa is an excellent system for understanding an individual’s antipredator behaviour in a group setting, as it eliminates the possibility of competition (for food and mates) confounding the expression of antipredator behaviours.

*Aedes aegypti* larvae and pupae are usually found in pools under high density (Ariyanto et al., 2020; Evans et al., 2023; Overgaard et al., 2017). While these immature stages do not exhibit coordinated movements, they are often aggregated near the walls of the containers or at the surface, resting together as they respire aerially. A dive by one larva or a pupa can make others dive in its vicinity, likely due to the water disturbance caused by the initial movement. A few studies indicate that these immature stages are aware of their conspecifics in water. For instance, Murthy et al., (2016) showed that *Aedes aegypti* larvae spend less time in the risky habitat when in a group than alone. Similarly, *Culex pipiens* pupae dive deep away from an attack if surrounded by fewer conspecifics, and the distance they flee decreases as the number of conspecifics increases (Rodríguez-Prieto et al., 2006). These studies highlight how the immature stages of mosquitoes can modulate their antipredator behaviour as per the conspecific density.

We have already established a clear behavioural carryover of predation-risk experience displayed by solitary pupa. When tested alone, pupae show an elaborate behaviour pattern in the absence and presence of an imminent threat that depends on their prior experience of predation risk (Rawat et al., 2024). Building on these findings, our current study aims to investigate whether similar behavioural carryovers occur in a group setting and compare it with our findings in solitary settings. To explore the expression of behavioural carryovers in a group setting, we reared larvae under conditions with and without predation risk. We then tested individual larvae at the pupal stage in group settings, focusing on key traits such as diving, space use, and overall activity levels in environments with and without an immediate threat. These traits were selected based on their relevance to survival under threat and their previously documented sensitivity to predation risk: diving is a key escape behaviour for mosquito larvae and pupae (Awasthi et al., 2012; Baglan et al., 2017; Romoser and Lucas, 1999); across diverse taxa, individuals adjust their spatial use and activity patterns in response to predation risk—for instance, reducing activity under immediate threat and seeking safer microhabitats (Abramsky et al., 1996; Blanchard et al., 2018; Ross et al., 2013). These behavioural traits were compared between risk-experienced and naive pupae. Hence, we compared the behaviour of experienced and naive pupae across two threat levels. Finally, we evaluated these results in light of our earlier findings on behavioural carryover in solitary settings to determine any consistencies or divergences in the patterns of risk-related behavioural carryover.

## Material and methods

The 2 x 2 experimental design involved exposing larvae to two distinct growth environments: one with predation cues and one without. Once the larvae developed into pupae, their behaviour—including dive frequency, dive depth, variation in dive depth, space use, and activity—was observed in two assay environments, with and without predation cues.

We used *Bradinopyga geminata* nymph as the common predator of *Aedes aegypti* larvae and pupae (Sharma et al., 2020; Venkatesh & Tyagi, 2013). We grew the first larval instars in brown trays (25 cm diameter and 5 cm height) with two litres of tap water and 0.15 g of food (dog biscuits and yeast in a 3:2 ratio). Forty-five larvae were maintained per tray. Half of the trays reared risk-experienced larvae with two dragonfly nymphs enclosed in 50 ml plastic containers; the other half reared predator-naive larvae. The larvae experienced the predation risk through visual and chemical cues but were not directly attacked. Just before pupation, we transferred the larvae to trays with no predation cues to maintain the naivete of the pupal stage.

The pupae were then recorded in two behavioural assay environments: under immediate threat (using water with predation cues) or no immediate threat environments (control tap water). This created four assay treatments– naive pupae in the absence of an immediate threat, naive pupae in the presence of an immediate threat, experienced pupae in the absence of an immediate threat, and experienced in the presence of an immediate threat. We recorded pupae in groups of five. Three individuals per group were randomly selected and followed for behavioural data on dive patterns, space use, and activity. The pupae were recorded for ten minutes in a 10 x 10 x 10 cm water column, of which the first three minutes were treated as an acclimation period.

We continuously followed the vertical movements of pupae where they left the surface and entered the water column, recording the deepest point they reached (measured at 2 cm resolution). These measures were used to calculate dive frequency, median dive depth, and coefficient of variation for dive depth to capture movement patterns and unpredictability. Space use was examined by following the location of pupae in the water column divided into the surface (breathing zone), middle, and bottom (the last 2cm, predator zone) every 10 seconds. These observations were used to calculate the proportion of time spent at each layer. Activity levels were determined by tracking transitions between passive and active states, calculating the proportion of time spent showing jerky muscular movements during the observation period. We extracted data from 48 individuals (3 x 16 groups) per assay treatment across two blocks of experiments in February and July 2022. Two observers cross-checked each data entry. Detailed experimental protocols are provided in Rawat et al., 2024.

### Statistical analyses

We analysed pupal behaviour data using mixed-effect models. Larval experience, pupal threat environment, experimental blocks, and the interaction between experience and threat environment were treated as fixed effects. Group IDs were included as a random effect, as each group contributed three individuals to the analysis. We used different mixed-effects models to test the influence of the predictor variables on various response variables (statistical tests with the R packages, residual distribution, link function and rationale are detailed in Table S1). We used a generalised mixed-effects model for dive frequency, and linear mixed-effects models for median dive depth and coefficient of variation in dive depth. We employed a multinomial mixed-effects model to quantify the effect of predictor variables on the relative proportion of time spent in different water columns by pupae. A beta mixed-effects regression model was used to measure the proportion of time spent in an active state. The robustness of the models was assessed by bootstrapping model confidence intervals. All statistical analyses were conducted using R version 4.2.3.

Likelihood ratio tests (LRT) and model confidence intervals were used to interpret the interaction term effects. With LRT *P* > 0.05 and 95% confidence intervals of coefficients overlapping zero, all the statistical models with the interaction term did not explain variation in the behavioural data better than those without the interaction term (Table S2, S3 and S4). This indicates that the interactive effect of larval experience and pupal threat environment did not explain the data better than their independent effects. Therefore, our interpretations are based on the models that include only the main effects.

## Results

### Carryover trends for dive pattern

Pupal diving behaviour was not discernibly affected by larval experience or immediate threat in the group setting (Table 1). Both naive and experienced pupae exhibited similar responses across conditions: dive frequency ranged from 5 to 7 dives per 7 minutes, median dive depth ranged from 7 cm to 8 cm, and variation in dive depth ranged from 30% to 40% of the mean (Fig 1). This pattern contrasts with the solitary setting, where experienced pupae demonstrated distinct behaviour compared to naive pupae. Specifically, experienced pupae exhibited higher dive frequency, more variable dive depths, and shallower dives in the absence of an immediate threat. They also altered their behaviour upon exposure to an immediate threat (Rawat et al., 2024).

**Fig 1:**
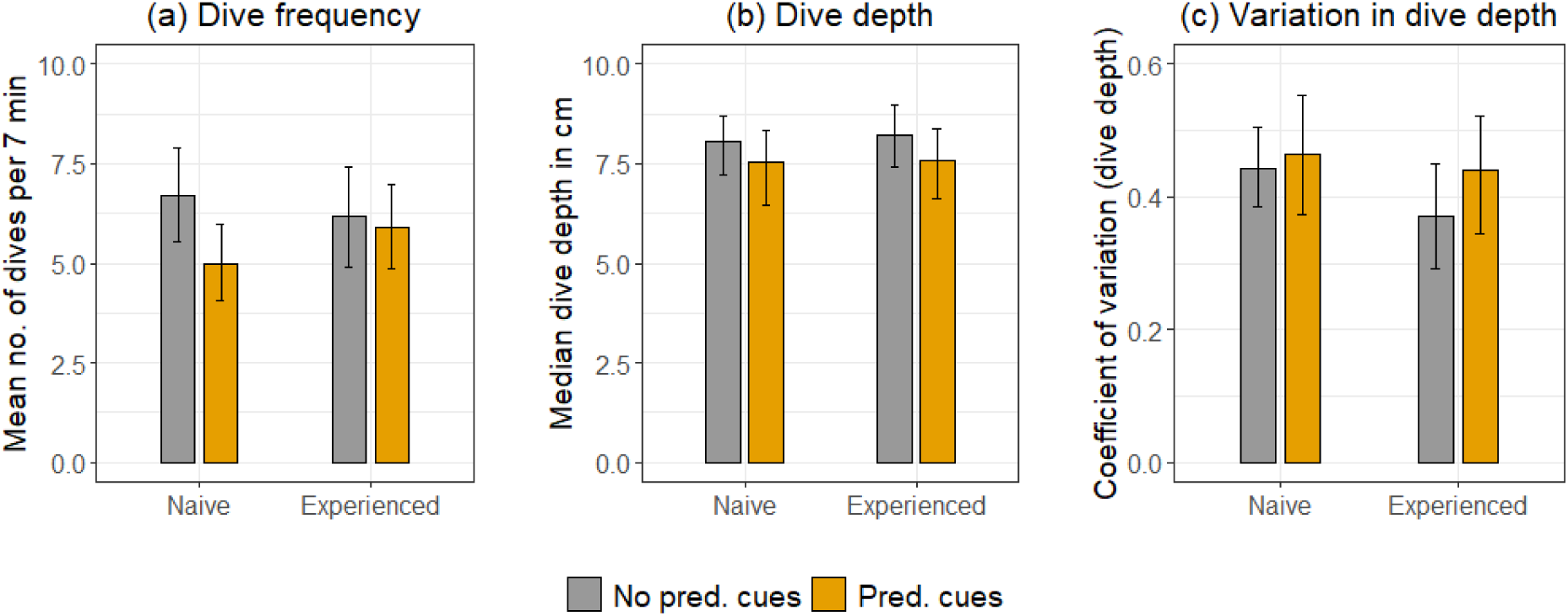
The bars indicate (a) mean dive frequency, (b) median dive depth, and (c) variation in dive depth for experienced pupae compared to naive pupae, both in the absence and presence of an immediate threat, in a group setting. Error bars represent 95% bootstrap confidence intervals. The colours represent the behavioural assay environment: grey indicates the absence of an immediate threat, while yellow indicates the presence of an immediate threat.

**Table 1:** Model estimates, 95% confidence intervals and P values from three main-effects models for response variables– dive frequency per 7 min, median dive depth and coefficient of variation of dive depth. We tested the effects of larval predation experience (experienced and naive) and pupal threat environment (absence and presence of predation cues) on the response variables using negative binomial GLMM for dive frequency and LMMs for dive depth and variation in dive depth.

|  | Dive frequency |  |  | Dive depth |  |  | Variation in dive depth |  |  |
| --- | --- | --- | --- | --- | --- | --- | --- | --- | --- |
|  | Estimate/<br>coefficient | 95%<br>Conf.<br>interval | P value | Estimate/<br>coefficient | 95%<br>Conf.<br>interval | P value | Estimate/<br>coefficient | 95%<br>Conf.<br>interval | P value |
| Intercept<br>(block1:naive:<br>control) | 1.782 | 1.587<br>1.978 | <0.0001 | 8.376 | 7.509<br>9.243 | <0.0001 | 0.372 | 0.283<br>0.462 | <0.0001 |
| block2 | 0.091 | -0.097<br>0.279 | 0.345 | -0.531 | -1.382<br>0.318 | 0.220 | 0.099 | 0.011<br>0.187 | 0.025 |
| experienced | 0.044 | -0.143<br>0.231 | 0.644 | 0.091 | -0.751<br>0.935 | 0.831 | -0.046 | -0.133<br>0.040 | 0.290 |
| cue | -0.170 | -0.357<br>0.017 | 0.075 | -0.570 | -1.413<br>0.272 | 0.184 | 0.045 | -0.041<br>0.132 | 0.304 |

### Carryover trends for space use pattern

Pupal space use pattern substantially varied with immediate threat environment but not larval experience. Both naive and experienced pupae moved from the middle water column toward the surface upon encountering predation cues (Table 2). Naive and experienced pupae decreased their time at the middle column by 7% to 13% and increased their time at the surface by 10% to 16% under the immediate threat environment. Meanwhile, 14% to 19% of the time was spent at the bottom, which did not change based on larval experience or the immediate threat environment (Fig 2). These results are in contrast to the solitary setting, where there was a small and variable effect of an immediate threat on space use.

**Fig 2:**
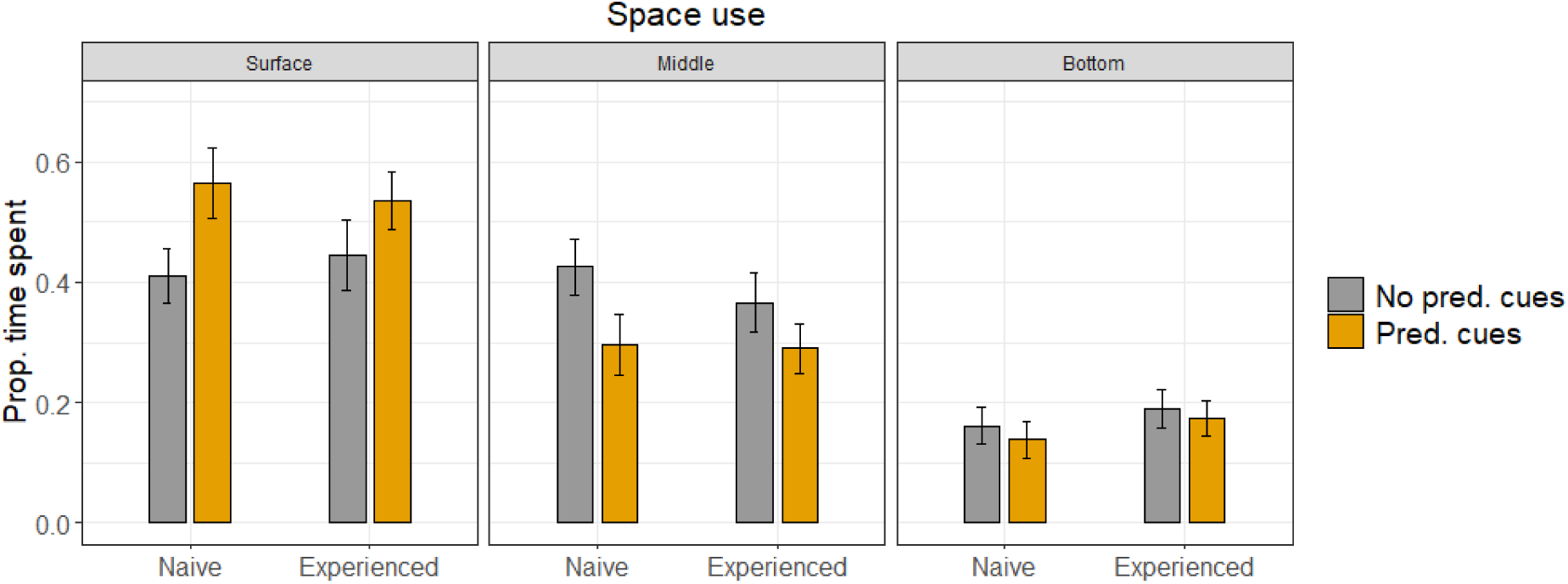
Comparison of carryover trends in space use patterns of naive and experienced pupae, focussing on the surface vs. middle water column. The bars represent the mean proportion of time spent at different water layers by experienced pupae compared to naive pupae, both in the absence and presence of an immediate threat, in a group setting. The error bars indicate the 95% bootstrap confidence intervals.

**Table 2:**
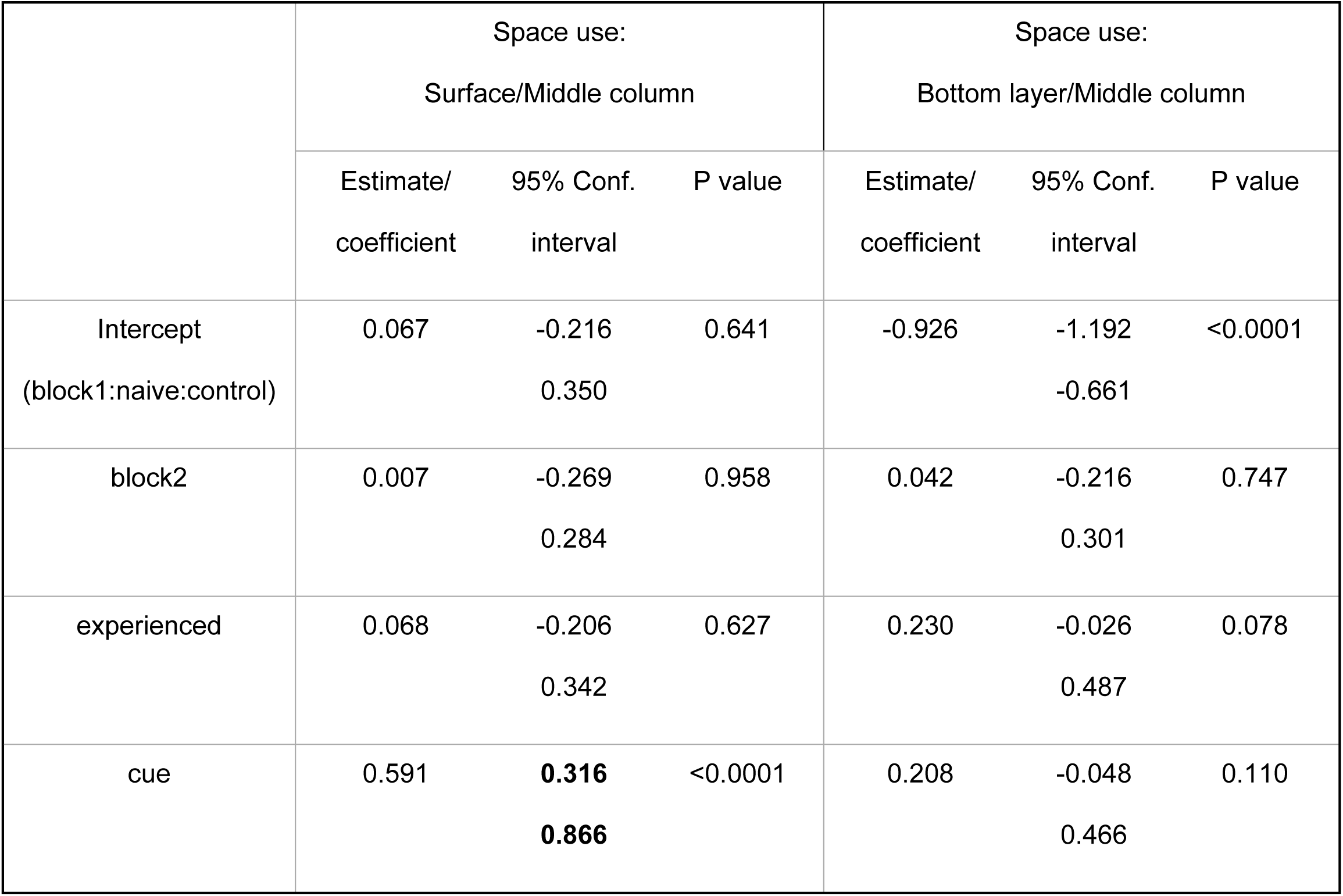
Model estimates, 95% confidence intervals and P values from main-effects multinomial GLMM used to test the effects of larval predation experience (experienced and naive) and pupal threat environment (absence and presence of predation cues) on the response variables– proportion time spent at the surface and bottom relative to the middle column. The detectable effects are indicated in bold.

|  | Space use:<br>Surface/Middle column |  |  | Space use:<br>Bottom layer/Middle column |  |  |
| --- | --- | --- | --- | --- | --- | --- |
|  | Estimate/<br>coefficient | 95% Conf.<br>interval | P value | Estimate/<br>coefficient | 95% Conf.<br>interval | P value |
| Intercept<br>(block1:naive:control) | 0.067 | -0.216<br>0.350 | 0.641 | -0.926 | -1.192<br>-0.661 | <0.0001 |
| block2 | 0.007 | -0.269<br>0.284 | 0.958 | 0.042 | -0.216<br>0.301 | 0.747 |
| experienced | 0.068 | -0.206<br>0.342 | 0.627 | 0.230 | -0.026<br>0.487 | 0.078 |
| cue | 0.591 | **0.316**<br>**0.866** | <0.0001 | 0.208 | -0.048<br>0.466 | 0.110 |

### Carryover trends for activity

Contrary to the finding that it had no effect in the solitary setting, the immediate threat environment affected the activity of both experienced and naive pupae in a group setting (Table 3). Experienced and naive pupae decreased the time spent being active on exposure to predation cues by 7% to 13% (Fig 3).

**Fig 3:**
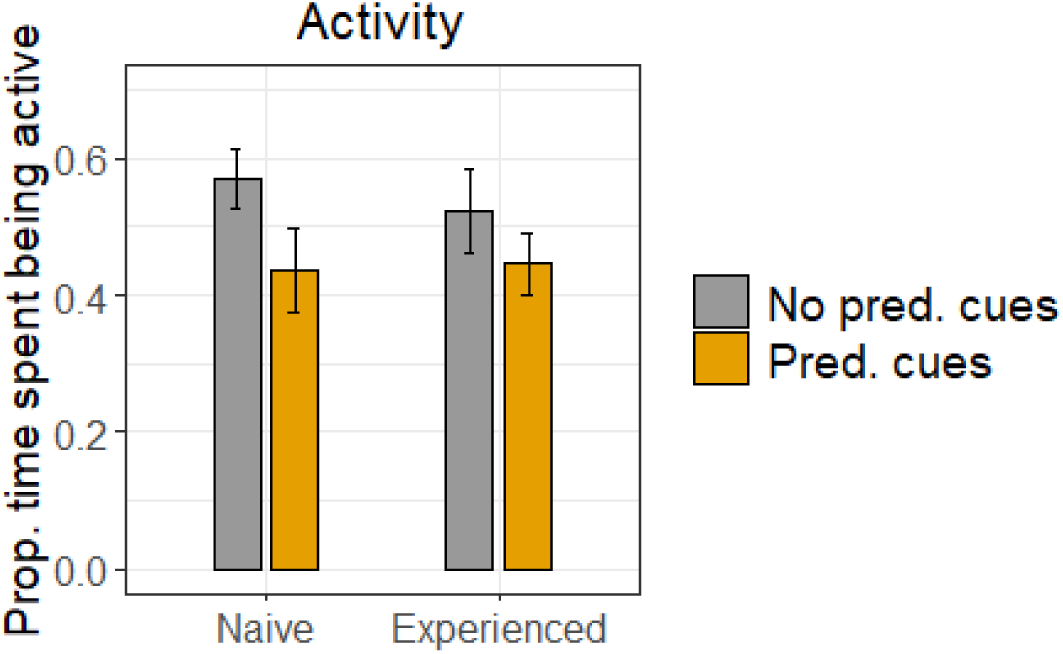
Comparison of carryover effects on the activity pattern in a group setting. The bars represent the mean proportion of time spent being active by experienced pupae compared to naive pupae, both in the absence and presence of an immediate threat, in a group setting. Error bars indicate 95% bootstrap confidence intervals.

**Table 3:** Model estimates, 95% confidence intervals and P values from the main-effects beta GLMM used to test the effects of larval predation experience (experienced and naive) and pupal threat environment (absence and presence of predation cues) on the response variables– proportion time spent active. The detectable effects are indicated in bold.

|  | Activity |  |  |
| --- | --- | --- | --- |
|  | Estimate/<br>coefficient | 95% Conf.<br>interval | P value |
| Intercept<br>(block1:naive:control) | 0.290 | 0.031 0.578 | 0.120 |
| block2 | -0.086 | -0.386 0.146 | 0.633 |
| experienced | -0.211 | -0.521 0.006 | 0.240 |
| cue | -0.405 | **-0.597 -0.068** | 0.024 |

## Discussion

The predation-risk experience of the larval stage did not influence the behaviour of the pupal stage in a group setting. In contrast to the solitary setting, where experienced pupae exhibited more frequent and shallower dives with greater variability in depth compared to naive pupae, the group setting eliminated these behavioural differences. This raises the question: why did experienced individuals cease to show the same responses in a group setting as they did alone? The answer likely lies in the different payoffs associated with behaviour in a group setting.

A group is an antipredator shield for an individual; therefore, the valuable behaviour of a solitary individual may add less value in protecting the individual from a predator in a group setting. In such instances, if the behaviour, such as diving, is energetically costly, (Lucas & Romoser, 2001), it is best avoided. Additionally, movement-related behaviour may also be costly as it can make a group more conspicuous to a predator; for example, Krause & Godin, 1995 showed that cichlid fish attacked shoals exhibiting more movement than groups that showed less movement. Sometimes, a group may interfere with specific movements; for example, grouped cowtail stingrays switch their escape angles to avoid collisions that reduce their escape speed from a predator (Semeniuk & Dill, 2004).

In addition to our result of no behavioural carryover of the past risk experience, we found a strong effect of the current threat environment on both naive and experienced pupae in group settings. Regardless of their previous risk experience, pupae changed their space use and activity patterns in the immediate threat environment. Naive and experienced pupae moved to the surface upon encountering predation cues and reduced their activity. The reduced activity is most likely due to spending more time at the surface where pupae rest. We came across a similar but weak and variable switch in space use by both naive and experienced pupae in solitary settings.

The substantial effect of the immediate threat environment on naive pupae underscores the significance of an innate pupal response against a potential threat, which is amplified in a group. Various social processes can explain this behaviour. Individuals respond to social information or cues to improve their foraging, mating and antipredator strategies (Danchin et al., 2004; Morand-Ferron et al., 2010). Under predation risk, naive individuals mimic conspecific behaviours and become more cohesive. For example, zebrafish are known to match behaviour by visual evaluation of their conspecifics that experience predation threat just (Oliveira et al., 2017), and white-breasted mesites increase cohesion following an alarm call (Gamero & Kappeler, 2015). Even though less explored, such trends are observed in visibly non-grouping animals (Tóth et al., 2020; Webster, 2023). For example, western spadefoot toad larvae mirror the behaviour of risk-exposed larvae (Caballero-Díaz et al., 2023), and wood crickets that observe the conspecifics demonstrating antipredator behaviour also show a hiding behaviour even in the absence of the demonstrators (Coolen et al., 2005).

The idea of social risk information transfer in the tiny pupal stage is compelling. However, it is important to rule out the possibility of individuals changing their behaviour in response to a threat based on the sheer number of conspecifics around them as it modifies their vulnerability. We can formally test the social transmission of information by seeding a behaviour in a group (Duboscq et al., 2016) and checking if the behaviour spreads in the group. The behaviour should be directed towards a cue that is not a natural predator to avoid stimulating naive individuals’ innate responses.

We observed that sociality governed individual behaviour in ways that downweight the value of prior experience. However, this does not negate the potential importance of past risk experience in group settings. Although we did not examine group-level behaviours and patterns in our study, it is known that certain group decisions, such as shoal cohesion in fish (Ioannou, 2017) and flock size in birds (Beauchamp, 2004), are often associated with high-risk environments.

## Conclusion

Our results highlight the importance of sociality in changing the payoffs of behavioural carryover and, arguably, the transmission of risk information even in non-grouping animals. They also emphasise the need to study seemingly simple life stages capable of making complex behavioural decisions in various contexts and understanding their associated payoffs.

## Acknowledgements

We express our gratitude to the Indian Institute of Science for providing institutional and logistical support. We sincerely appreciate the assistance in colony maintenance and knowledge sharing from Kanchana Gaonkar, Manvi Sharma, Karthikeyan Chandrasegaran, and Akshay P. Special thanks go to our project assistants: Akshaye Anand Bhambore, Gokul Bhaskaran, Shubhada Shirish More and Navina Mable Francis, as well as our dedicated team of interns: Subhiksha M Bharadwaj, Aranya Dhibar, Ashmita Baruah, Reva T, Jitty Alin Jacob, Hareendran K M, Nidhi Yadav, Ishika and Sukanya, for their invaluable support during experiments, troubleshooting, data extraction and cross-checking. We also extend our thanks to our friends for their essential help during fieldwork. We are grateful for fellowship support from the Ministry of Education of India and grant support from the Department of Science and Technology-Science and Engineering Research Board (DST-SERB) through the Scientific and Useful Profound Research Advancement (SUPRA) grant, SPR/2020/000384. Additionally, we acknowledge funding from DST-FIST [Fund for Improvement of S&T Infrastructure; sanction number SR/FST/LSII-025/2009(C)], the DBT-IISc Partnership Program (Phase II; Department of Biotechnology, Ministry of Science and Technology and Indian Institute of Science, India), CSIR (Council of Scientific and Industrial Research) 37(1636)/14/EMR-II, and the Indian Institute of Science. Finally, we acknowledge the approval from the Institutional Animal Ethics Committee (constituted by the Indian Institute of Science, CAF/Ethics/835/2021) for breeding mosquitoes in the lab and conducting experiments with live insects.

## References

Abramsky, Z., Strauss, E., Subach, A., Riechman, A., & Kotler, B. P. (1996). The effect of barn owls (*Tyto alba*) on the activity and microhabitat selection of *Gerbillus allenbyi* and *G. pyramidum*. Oecologia, 105, 313–319.

Ariyanto, A., Ibrahim, E., Syahribulan, S., Ishak, H., Syamsuar, S., & Djajakusli, R. (2020). Density of *Aedes aegypti* larvae based on knowledge, attitude, and action to eradicate mosquito nest in Daya market of Makassar city. Journal of Asian Multicultural Research for Medical and Health Science Study, 1(2), Article 2. 10.47616/jamrmhss.v1i2.52

Awasthi, A. K., Wu, C. H., & Hwang, J. S. (2012). Diving as an anti-predator behavior in mosquito pupae. Zoological Studies, 51(8), 1225–1234.

Baglan, H., Lazzari, C., & Guerrieri, F. (2017). Learning in mosquito larvae (*Aedes aegypti)*: Habituation to a visual danger signal. Journal of insect physiology, 98, 160–166. 10.1016/j.jinsphys.2017.01.001

Beauchamp, G. (2004). Reduced flocking by birds on islands with relaxed predation. Proceedings of the Royal Society of London. Series B: Biological Sciences, 271(1543), 1039–1042. 10.1098/rspb.2004.2703

Blanchard, P., Lauzeral, C., Chamaillé-Jammes, S., Brunet, C., Lec’hvien, A., Péron, G., & Pontier, D. (2018). Coping with change in predation risk across space and time through complementary behavioral responses. BMC ecology, 18. 10.1186/s12898-018-0215-7

Brass, K. E., Herndon, N., Gardner, S. A., Grindstaff, J. L., & Campbell, P. (2020). Intergenerational effects of paternal predator cue exposure on behavior, stress reactivity, and neural gene expression. Hormones and Behavior, 124, 104806. 10.1016/j.yhbeh.2020.104806

Brown, G. E., & Godin, J.-G. J. (2023). Ecological uncertainty and antipredator behaviour: An integrative perspective. Frontiers in Ethology, 2. https://www.frontiersin.org/articles/10.3389/fetho.2023.1238167

Caballero-Díaz, C., Arribas, R., & Polo-Cavia, N. (2023). Assessment of predation risk through conspecific cues by anuran larvae. Animal Cognition, 26(4), 1431–1441. 10.1007/s10071-023-01793-y

Coolen, I., Dangles, O., & Casas, J. (2005). Social learning in noncolonial insects? Current Biology, 15(21), 1931–1935. 10.1016/j.cub.2005.09.015

Couzin, I. D., & Krause, J. (2003). Self-organization and collective behavior in vertebrates. In Slater, P., Rosenblatt, J., Snowdon, C., Roper, T. (Eds.), Advances in the Study of Behavior (Vol. 32, pp. 1–75). Elsevier. 10.1016/S0065-3454(03)01001-5

da Cunha, F. C. R., Fontenelle, J. C. R., & Griesser, M. (2017). Predation risk drives the expression of mobbing across bird species. Behavioral Ecology, 28(6), 1517– 1523. 10.1093/beheco/arx111

Danchin, É., Giraldeau, L.-A., Valone, T. J., & Wagner, R. H. (2004). Public information: From nosy neighbors to cultural evolution. Science, 305(5683), 487–491. 10.1126/science.1098254

Morand-Ferron, J., Doligez, B., Dall, S. R. X., & Reader, S. M. (2010). Social information use. Encyclopedia of Animal Behavior, 3. 242–250. 10.1016/B978-0-08-045337-8.00281-3

Downes, S., & Hoefer, A. M. (2004). Antipredatory behaviour in lizards: Interactions between group size and predation risk. Animal Behaviour, 67(3), 485–492. 10.1016/j.anbehav.2003.05.010

Duboscq, J., Romano, V., MacIntosh, A., & Sueur, C. (2016). Social information transmission in animals: Lessons from studies of diffusion. Frontiers in Psychology, 7, 1147. 10.3389/fpsyg.2016.01147

Evans, K. G., Neale, Z. R., Holly, B., Canizela, C. C., & Juliano, S. A. (2023). Survival-larval density relationships in the field and their implications for control of container-dwelling *Aedes* mosquitoes. Insects, 14(1), Article 1. 10.3390/insects14010017

Ezenwa, V. O., Ghai, R. R., McKay, A. F., & Williams, A. E. (2016). Group living and pathogen infection revisited. Current Opinion in Behavioral Sciences, 12, 66–72. 10.1016/j.cobeha.2016.09.006

Gamero, A., & Kappeler, P. M. (2015). Always together: Mate guarding or predator avoidance as determinants of group cohesion in white-breasted mesites? Journal of Avian Biology, 46(4), 378–384.

Garcia, T. S., Bredeweg, E. M., Urbina, J., & Ferrari, M. C. O. (2019). Evaluating adaptive, carry-over, and plastic antipredator responses across a temporal gradient in Pacific chorus frogs. Ecology, 100(11), e02825. 10.1002/ecy.2825

Groenewoud, F., Kingma, S. A., Bebbington, K., Richardson, D. S., & Komdeur, J. (2019). Experimentally induced antipredator responses are mediated by social and environmental factors. Behavioral Ecology, 30(4), 986–992. 10.1093/beheco/arz039

Ioannou, C. (2017). Grouping and predation. In Shackelford, T. K. & Weekes-Shackelford, V. A. (Eds.), Encyclopedia of Evolutionary Psychological Science (pp. 1–6). Springer International Publishing. 10.1007/978-3-319-16999-6_2699-1

Krause, J., & Godin, J.-G. J. (1995). Predator preferences for attacking particular prey group sizes: Consequences for predator hunting success and prey predation risk. Animal Behaviour, 50(2), 465–473. 10.1006/anbe.1995.0260

Landry, F., & Li, M. F. (2022). Costs of group living. In Vonk, J. & Shackelford, T. K. (Eds.), Encyclopedia of Animal Cognition and Behavior (pp. 1744–1750). Springer International Publishing. 10.1007/978-3-319-47829-6_295-1

Lehtonen, J., & Jaatinen, K. (2016). Safety in numbers: The dilution effect and other drivers of group life in the face of danger. Behavioral Ecology and Sociobiology, 70(4), 449–458. 10.1007/s00265-016-2075-5

Lucas, E. A., & Romoser, W. S. (2001). The energetic costs of diving in *Aedes aegypti* and *Aedes albopictus* pupae. Journal of the American Mosquito Control Association, 17(1), 56–60.

Murthy, A., Sharma, M., Amith-Kumar, U. R., & Isvaran, K. (2016). Groups constrain the use of risky habitat by individuals: A new cost to sociality? Animal Behaviour, 113, 167–175. 10.1016/j.anbehav.2015.12.027

Oliveira, T. A., Idalencio, R., Kalichak, F., dos Santos Rosa, J. G., Koakoski, G., de Abreu, M. S., Giacomini, A. C. V., Gusso, D., Rosemberg, D. B., Barreto, R. E., & Barcellos, L. J. G. (2017). Stress responses to conspecific visual cues of predation risk in zebrafish. PeerJ, 5, e3739. 10.7717/peerj.3739

Overgaard, H. J., Olano, V. A., Jaramillo, J. F., Matiz, M. I., Sarmiento, D., Stenström, T. A., & Alexander, N. (2017). A cross-sectional survey of *Aedes aegypti* immature abundance in urban and rural household containers in central Colombia. Parasites & Vectors, 10(1), 356. 10.1186/s13071-017-2295-1

Rawat, K., Bhambore, A. A., & Isvaran, K. (2024). Don’t leave the past behind: How larval experience shapes pupal antipredator response in Aedes aegypti (p. 2024.02.02.578532). bioRxiv. 10.1101/2024.02.02.578532

Rodríguez-Prieto, I., Fernández-Juricic, E., & Martín, J. (2006). Anti-predator behavioral responses of mosquito pupae to aerial predation risk. Journal of Insect Behavior, 19(3), 373–381. 10.1007/s10905-006-9033-4

Romoser, W. S., & Lucas, E. A. (1999). Buoyancy and diving behavior in mosquito pupae. Journal of the American Mosquito Control Association, 15(2), 194–199.

Ross, J., Hearn, A. J., Johnson, P. J., & Macdonald, D. W. (2013). Activity patterns and temporal avoidance by prey in response to Sunda clouded leopard predation risk. Journal of Zoology, 290(2), 96–106.

Rubenstein, D. (1978). On predation, competition, and the advantages of group Living. In Bateson, P. P. G., Klopfer, P. H. (Eds.), Perspectives in Ethology (Vol. 3, pp. 205–231). 10.1007/978-1-4684-2901-5_9

Semeniuk, C. A. D., & Dill, L. M. (2005). Cost/benefit analysis of group and solitary resting in the cowtail stingray, *Pastinachus sephen*. Behavioral Ecology, 16(2), 417–426. 10.1093/beheco/ari005

Sharma, M., Quader, S., Guttal, V., & Isvaran, K. (2020). The enemy of my enemy: Multiple interacting selection pressures lead to unexpected anti-predator responses. Oecologia, 192(1), 1–12. 10.1007/s00442-019-04552-4

Tóth, Z., Jaloveczki, B., & Tarján, G. (2020). Diffusion of social information in non-grouping animals. Frontiers in Ecology and Evolution, 8. 10.3389/fevo.2020.586058

Venkatesh, A., & Tyagi, B. (2013). Predatory potential of *Bradinopyga geminata* and *Ceriagrion coromandelianum* larvae on dengue vector *Aedes aegypti* under controlled conditions (Anisoptera: Libellulidae; Zygoptera: Coenagrionidae; Diptera: Culicidae). Odonatologica, 42(2), 139–149.

Wang, Y., Fu, S.-J., & Fu, C. (2019). Behavioral adjustments to prior predation experience and food deprivation of a common cyprinid fish species vary between singletons and a group. PeerJ, 7, e7236. 10.7717/peerj.7236

Webster, M. M. (2023). Social learning in non-grouping animals. Biological Reviews, 98(4), 1329–1344. 10.1111/brv.12954

West, R., Letnic, M., Blumstein, D. T., & Moseby, K. E. (2018). Predator exposure improves anti-predator responses in a threatened mammal. Journal of Applied Ecology, 55(1), 147–156. 10.1111/1365-2664.12947

Wirsing, A. J., Heithaus, M. R., Brown, J. S., Kotler, B. P., & Schmitz, O. J. (2021). The context dependence of non-consumptive predator effects. Ecology Letters, 24(1), 113–129. 10.1111/ele.13614

