## Supplementary material for "Peers over the past: Prior predation-risk experience does not dictate antipredator responses of individuals in groups": Table S2,

| Behavioural trait | Statistical test | R package used | Distribution | Link function | Rationale based on the nature of the response variable |
|---|---|---|---|---|---|
| Dive frequency | Negative binomial GLMM | glmmTMB | Negative Binomial distribution | Log link function | Overdispersed count data |
| Dive depth | Linear mixed-effects model | glmmTMB | Normal (Gaussian) distribution | Identity link function | Continuous variable |
| Variation in dive depth | Linear mixed-effects model | glmmTMB | Normal (Gaussian) distribution | Identity link function | Continuous variable |
| Space use | Multinomial mixed-effects model | mclogit | Multinomial distribution | Logit link function | Multiple discrete outcomes in no specific order |
| Activity | Beta regression | glmmTMB | Beta distribution | Logit link function | Proportion data |

Table S1: Details of statistical models used for different behavioural traits, along with the rationales based on the nature of the response variables

| | Dive frequency | | Dive depth | | Variation in dive depth | |
|---|---|---|---|---|---|---|
| | **Estimate** | **95% Conf. interval** | **Estimate** | **95% Conf. interval** | **Estimate** | **95% Conf. interval** |
| **Intercept (block1:naive:control)** | 1.843 | 1.633 2.054 | 8.341 | 7.385 9.297 | 0.386 | 0.287 0.484 |
| **block2** | 0.095 | -0.090 0.281 | -0.532 | -1.383 0.318 | 0.100 | 0.012 0.187 |
| **experienced** | -0.078 | -0.335 0.177 | 0.163 | -1.012 1.340 | -0.074 | -0.196 0.047 |
| **cue** | -0.298 | -0.561 -0.036 | -0.496 | -1.692 0.700 | 0.016 | -0.106 0.140 |
| **experienced:cue** | 0.252 | **-0.115 0.621** | -0.148 | **-1.834 1.538** | 0.056 | **-0.117 0.230** |

Table S2: Model estimates and 95% confidence intervals from three models testing interaction effects on the response variables—dive frequency per 7 minutes, median dive depth, and the coefficient of variation of dive depth—are presented. We examined the interactive effect of larval predation-risk experience and pupal threat environment on dive frequency using a negative binomial GLMM, and on median dive depth and variation in dive depth using linear mixed models (LMMs). Interaction terms with confidence intervals overlapping zero, indicating non-significance, are highlighted in bold.

| | Space use: Surface/Middle column | | Space use: Bottom layer/Middle column | | Activity | |
|---|---|---|---|---|---|---|
| | **Estimate** | **95% Conf. interval** | **Estimate** | **95% Conf. interval** | **Estimate** | **95% Conf. interval** |
| **Intercept (block1:naive:control)** | -0.019 | -0.329 0.291 | -0.985 | -1.278 -0.692 | 0.328 | -0.079 0.736 |
| **block2** | 0.006 | -0.267 0.279 | 0.042 | -0.216 0.300 | -0.086 | -0.443 0.269 |
| **experienced** | 0.241 | -0.141 0.624 | 0.347 | -0.010 0.705 | -0.286 | -0.786 0.214 |
| **cue** | 0.766 | 0.382 1.151 | 0.330 | -0.035 0.696 | -0.479 | -0.981 0.021 |
| **experienced:cue** | -0.350 | **-0.893 0.191** | -0.241 | **-0.753 0.270** | 0.148 | **-0.559 0.855** |

Table S3: Model estimates with 95% confidence intervals from two models: a multinomial GLMM with response variables as proportions of time spent at the surface and bottom layers relative to the middle layer; a beta GLMM with ‘proportion time spent being active’ as the response variable. We estimated the interactive effect of the larval predation-risk experience and pupal threat environment on the response variables. The interaction-term confidence intervals, overlapping zero, are indicated in bold.

| Trait | Likelihood ratio (chisq) | Degrees of freedom | P value |
|---|---|---|---|
| Dive frequency | 1.769 | 1 | 0.183 |
| Dive depth | 0.029 | 1 | 0.863 |
| Variation in dive depth | 0.406 | 1 | 0.524 |
| Space use | 1.743 | 2 | 0.418 |
| Activity | 0.274 | 1 | 0.600 |

Table S3: Likelihood ratio tests comparing the interaction model and the main-effects model for all the behavioural traits. We assessed the interaction term, ‘Larval growth env:Behavioural assay env’, by comparing the models with and without the interaction term. The interaction-effect models did not explain the data better than the main-effects models.
